# *De novo* assembly of the selfish *t* supergene reveals a deleterious evolutionary trajectory

**DOI:** 10.1101/2024.09.15.613113

**Authors:** Jan-Niklas Runge, Kristian Ullrich, Anna K. Lindholm

## Abstract

Supergenes are linked clusters of DNA that are transmitted together due to rare or absent recombination. They undergo co-adaptation, allowing evolution to work on several genes to refine complex phenotypes, giving supergenes a competitive edge. Yet, due to their lack of recombination, supergenes are susceptible to deterioration as they cannot efficiently purge deleterious DNA. Few examples outside of sex chromosomes have been available for study. Here, we present the first assembly of the *t* haplotype, a 33.4 Mb supergene in house mice that ‘selfishly’ transmits itself at non-Mendelian frequencies. We characterize the four large non-overlapping inversions that make up the *t* haplotype. We compare in a *t*/*t* individual two different *t* variants with different recessive lethal phenotypes (age at death). Despite that difference, they differ much less from each other than the rest of the chromosome. However, the differences that they have were much more likely to be deleterious than the differences between the two variants of the rest of the chromosome. We interpret this marked difference as evidence of the accumulation of deleterious variants, a hallmark of deterioration. The *t* region of chromosome 17 is more distant to the reference than the rest of the chromosome, and has a higher fraction of impactful differences here as well. Thus, we conclude that the *t* appears as a quickly spreading and deteriorating selfish supergene, a rare example of Muller’s ratchet outside of sex chromosomes. Furthermore, we aim for our assembly to provide a resource for comparative work on the *t* haplotype, such as its evolutionary history.

---

Complex, adaptive phenotypes often require multiple genes to function, but their evolution is constrained by recombination, which breaks up linkage of alleles. Inversions can ensure that such combinations of alleles remain together and are inherited as a unit, called a supergene. Phenotypes such as social organization and mating types, but also sexual differentiation and selfish transmission are built on this mechanism, but the reduction in recombination necessary for joint transmission can also decrease the supergene’s fitness.

The reduced recombination of supergenes is expected to result in their deterioration through accumulation of deleterious mutations and expansion of repetitive elements. This is because selection can no longer work on deleterious content within the supergene, but instead works on the supergene as a whole. When coupled with low or intermediate frequencies of supergenes, processes such as Muller’s ratchet and background selection (Charlesworth and Charlesworth 2000) are expected to lead to deterioration. In the process of deterioration, supergenes can accumulate repetitive DNA, which can increase the size of the supergene as in *Sb* in fire ants *Solenopsis invicta* (Stolle et al. 2019), but can ultimately also lead to its shortening, like in the mammalian *Y*, accompanied by gene loss (Graves 2006). Patterns can also be more complicated, such as in *Papilio* butterflies where the mimicry supergene locus has increased repetitive elements, both in inverted and non-inverted species (Komata et al. 2022). In addition to an increase in repetitive DNA, supergene deterioration can also lead to relative increases in non-synonymous mutations, evidence of less efficient purifying selection (Svedberg et al. 2018). However, both young and old supergenes can be found to be unassociated with an increase in repetitive elements or a great accumulation of deleterious mutations (Hill et al. 2022; Stenløkk et al. 2022). While young supergenes may still degenerate, it is unclear how degeneration is averted in cases of old supergenes.

The *t* haplotype is an autosomal male meiotic driver in house mice that is an old supergene, approximately 1-3 million years old (Morita et al. 1992; Hammer and Silver 1993). It is about 35 megabases (Kelemen and Vicoso 2018), a third of chromosome 17, in size, and is linked by four non-overlapping inversions (Herrmann et al. 1986; Hammer et al. 1989; Howard et al. 1990; Sugimoto 2014). The *t* is widespread in nature, but overall low in frequency, due to natural selection from deleterious, recessive alleles and sexual selection from poor sperm competitive ability (Dunn and Gluecksohn-Schoenheimer 1943; Sutter and Lindholm 2015; Manser et al. 2020). It is so far not known to what extent variants of the *t* differ from one another (Ardlie and Silver 1998; Manser et al. 2011; Sugimoto 2014). Furthermore, the actual content of the *t* and the breakpoints of the inversions have so far mostly been studied by comparison to the reference house mouse genome and pre-NGS mapping methods, except for recent short-read-based genomics and transcriptomics work (Kelemen and Vicoso 2018; Lindholm et al. 2019; Kelemen et al. 2022), despite considerable differences between *t* and other variants of chromosome 17. Here, we provide the first de novo assembly of the *t* haplotype, and analyze its contents.

## Results

### The *t*-carrying chromosome 17

By aligning contigs to the reference genome, we found five contigs that together constitute the *t*-carrying chromosome 17. To infer the orientation and order of the chromosome 17 contigs independently of the reference genome, we inspected paired-end reads that mapped onto different contigs and PacBio reads that mapped similarly well onto ends of different contigs. We find that both approaches generally agree on the orientation between contigs 1-2 (3 paired-end reads; 60 long reads), 2-3 (3 paired-end reads; 43 long reads), 3-4 (20 PE reads, 594 long reads), and 4-5 (5 PE, 98 LR). Conflicting evidence did not go in a clearly different direction (**SI Table 1** & **SI Figure 1**), for example there was more longread evidence in favor of a flipped contig 2 into flipped contig 1 order (rather than flipped 1 to flipped 2), but no short reads supporting that and the order we arrived at is more parsimonious given the discovered inversion breakpoints within contigs (**Figure 1 A**) and the connections to the other contigs.

**Figure 1:**
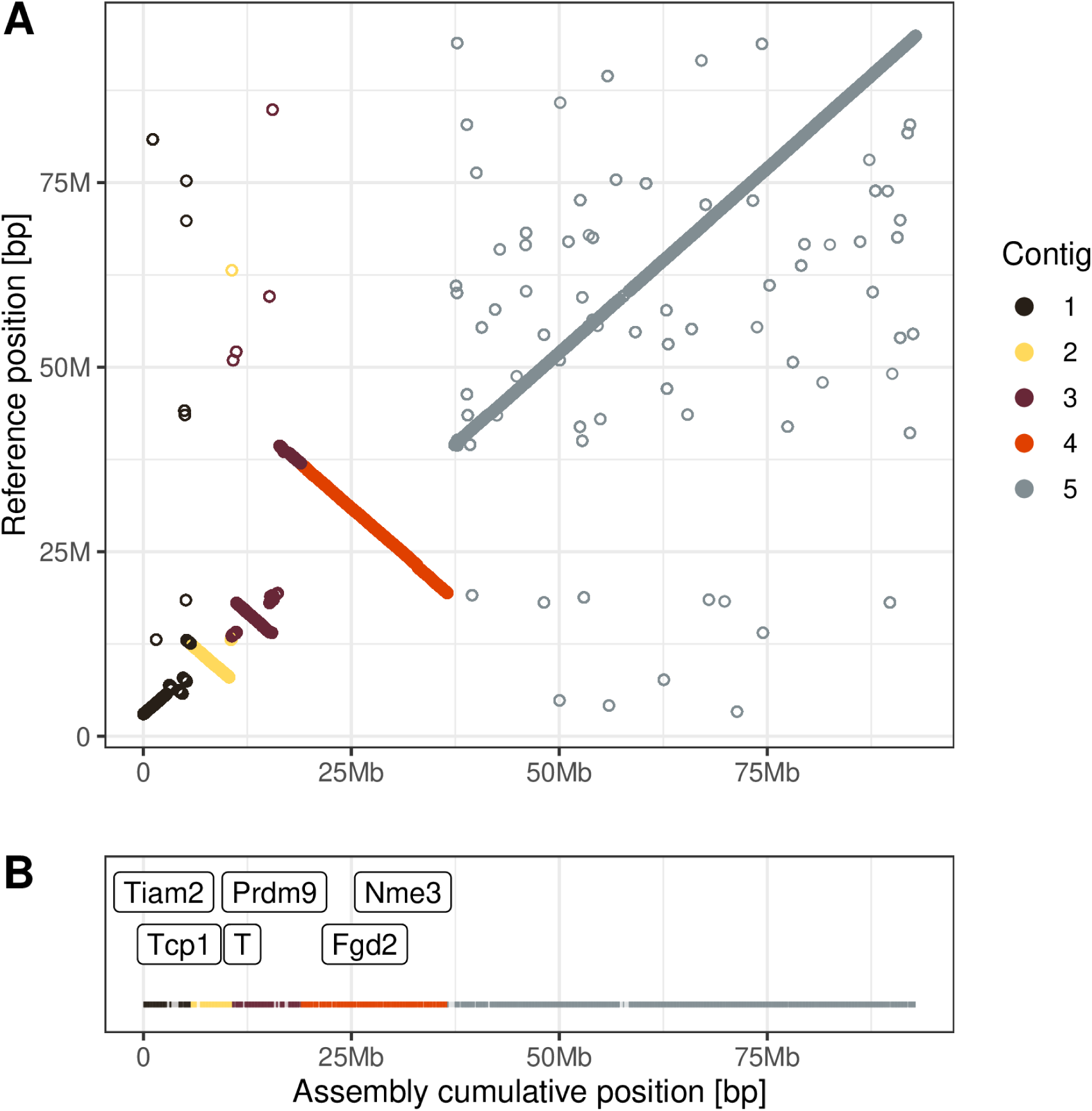
**A)** Dotplot of alignments between *t^wIll^/t^wGrabs^* chromosome 17 and reference house mouse chromosome 17. **B)** Indication of regions with missing bases via decreased visibility. Position of a selection of *t* genes indicated.

Mapping the concatenated assembly with the derived orientations of the contig against the reference reveals a pattern of four major inversions within the *t* contigs (**Figure 1 A**). Using the variant detection tool *SYRI* (Goel et al. 2019), we inferred the breakpoints of these major and some minor inversions (**Table 1**). Seven out of eight major inversion breakpoints are covered by our assembly **Figure 1 A**. Since the *t* is defined as a supergene with reduced recombination due to inversions (without known coordinates) (Howard et al. 1990; Sugimoto 2014), we henceforth refer to the region from the start of the first major inversion (∼ at 5.8 Mb on the reference) to the end of the last major inversion (∼ 39.3 Mb) as the *t* region on our assembly (33.4 Mb) and the rest as the *w* region of our assembly. This *t* region in our assembly is 99.5% of the size of the reference region in those borders. The remaining *w*region is 96.8% as large as the remaining reference region. Thus, our assembly likely covers close to the entire chromosome. However, our assembly includes regions with unknown bases (10.6% for *t* and 4.6% for *w* regions, **1 B**), likely due to insufficient coverage, which is important to note regarding our results, for example in gene annotations.

**Table 1:** Overview of the detected inversions in our chromosome 17 assembly, as well as their coordinates on the *GRCm38.p6* reference and our assembly.

| Name | Length | Begin Reference | End Reference | Start Assembly | End Assembly |
| --- | --- | --- | --- | --- | --- |
| Inv1 | 1152298 | 5766767 | 6919065 | 3123746 | 4711557 |
| Inv2 | 5029646 | 7987215 | 13016861 | 5200654 | 10320713 |
|  | 53998 | 13687929 | 13741927 | 10710468 | 10769301 |
|  | 12976 | 13987100 | 14000076 | 11034555 | 11047334 |
| Inv3 | 3948768 | 14102739 | 18051507 | 11205098 | 15161795 |
| Inv4 | 19919868 | 19407603 | 39327471 | 16396378 | 36534313 |
|  | 648 | 63755209 | 63755857 | 61804226 | 61804874 |
|  | 240 | 66460437 | 66460677 | 64523125 | 64523367 |
|  | 493 | 68453688 | 68454181 | 66537462 | 66537957 |
|  | 226 | 69915121 | 69915347 | 67984169 | 67984394 |
|  | 2298 | 82649353 | 82651651 | 80668583 | 80670934 |
|  | 459 | 90716921 | 90717380 | 88698027 | 88698516 |
|  | 479 | 91555764 | 91556243 | 89528087 | 89528569 |

### Comparison to known inversion coordinates

The 33.4 Mb size of the *t* region in our assembly, as well as its 5.8 Mb to 39.3 Mb coordinates when mapped against the house mouse reference genome are very similar to the region of increased *t*/*w* heterozygosity (mapped against the reference chromosome 17) described by Kelemen and Vicoso (2018) at 5 Mb to 40 Mb. However, the region between inversion 3 and 4 appears much smaller in our assembly. Nonetheless, we also find a clear size difference between inversion 4, inversions 2 and 3, and inversion 1, in contrast to how earlier work using pre-NGS methodologies described and visualized the inversions as two large (inv2 and inv4) and two small (inv1 and inv3) inversion (Sugimoto 2014).

### Annotation

We annotated the chromosome 17 assembly using *GeMoMa* and detected sequences of 749 unique genes in the *t* region, with 77% of them also found on the reference chromosome 17 and unplaced chromosome 17 contigs (compared to 598 and 59% for the *w* region, see *Data availability*).

Five out of eight canonical *t*-related genes were found in the *t* inversions **Figure 1 B**: *T* (Inv2 (Herrmann et al. 1986)), *Tcp1* (Inv2 (Hammer et al. 1989)), *Prdm9* (Inv3 (Kono et al. 2014)), *Fgd2* (Inv4 (Bauer et al. 2007)), and *Nme3* (Inv4 (Bauer et al. 2012)). One, *Tiam2* (Charron et al. 2019), a candidate distorter in the first inversion, was found well outside the first inversion (between the centromere and first inversion breakpoint), so by our definition outside of *t*. Two genes known to be critical for the *t*’s transmission distortion, *Tagap* and *Smok*, were not found (Herrmann and Bauer 2012). All 41 detected histocompatibility 2 complex (*H-2* or *MHC*) genes were found in the fourth *t* inversion (Sugimoto 2014).

### Differences between the two *t* variants

The *t* assembly is based on an offspring of parents from two wild strains (ILL and GRABS) that do not interbreed, which each carry one *t* variant (*t^wIll^* or *t^wGrabs^*), which differ at least in their recessive deleterious alleles (see Methods). To discover the degree to which they differ, we analyzed the base differences by mapping Illumina reads of pooled *t^wIll^/t^wIll^* homozygous embryos from one strain, as well as the most confident heterozygous SNP calls made by mapping the PacBio reads (*t^wIll^/t^wGrabs^*) back to the polished assembly, comparing the *t* and *w* regions of chromosome 17, with *w* serving as a baseline for the difference between the two strains.

51% more bases differed between our assembly and *t^wIll^/t^wIll^*short reads in the *w* region than in the *t* region, implying a larger difference between *w^wIll^* and *w^wGrabs^* than *t^wIll^* and *t^wGrabs^* (**Figure 2 A-B**). Using the heterozygosity of *t^wIll^/t^wGrabs^*long reads, the difference rose to 217% (**Figure 2 C-D**). Combined, the fraction of heterozygous SNPs per base for the *t* region was between 2*^−^*^4^ and 4*^−^*^4^, (long- and short-read based, respectively) and between 5*^−^*^4^ and 7*^−^*^4^ for the *w* region. Both of these numbers are more than halved from population-wide estimates of nucleotide diversity for wild house mice (between 2*^−^*^3^ for German and 3*^−^*^3^ for Iranian samples; Harr et al. (2016))

**Figure 2:**
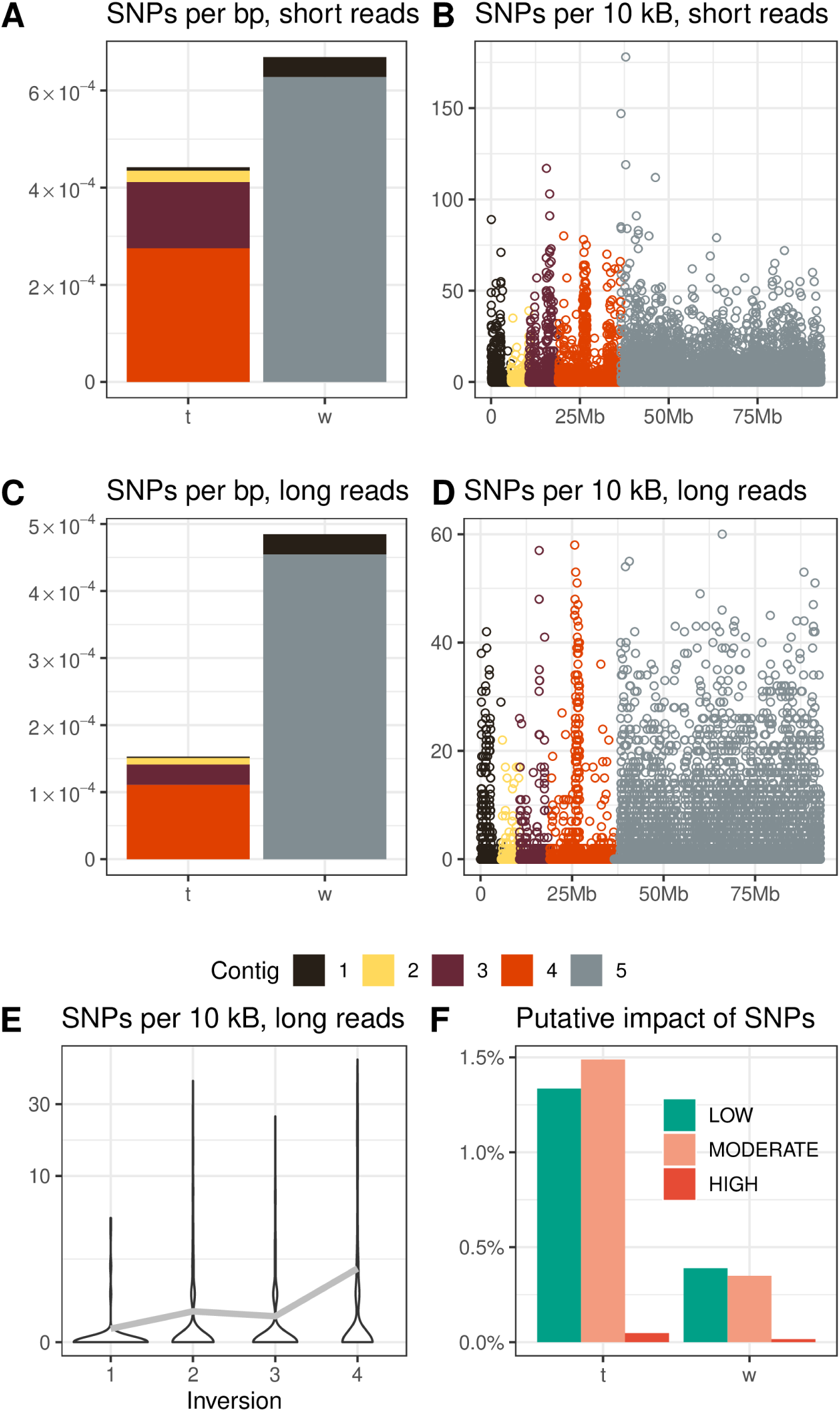
Heterozygous single nucleotide polymorphisms (SNPs) between GRABS and ILL per non-N base pair in the *t* and *w* regions of chromosome 17. **A)** SNPs per base pair based on *Pilon* polishing using *t^wIll^/t^wIll^*short reads. **B)** SNPs per 10 kB window based on *Pilon* polishing using *t^wIll^/t^wIll^*short reads. **C)** SNPs per base pair based on heterozygous long reads. **D)** SNPs per 10 kB window based on heterozygous long reads. **E)** Violin plots of SNPs per 10 kB within each inversion. Grey line indicates mean value per inversion. **F)** *SnpEff* estimated impact of heterozygous SNPs (excluding the lowest possible (non-)impact) in *t* and *w* regions in percent of variants.

We aggregated the number of *t* heterozygous SNPs per 10 kB, normalized by the number of unknown bases in each window, into the four major inversions (named as in Kelemen and Vicoso (2018)) and found large differences between them (**Figure 2 E**). Unsurprisingly, the large heterozygosity peak in inversion four means that this inversion is the most diverse within *t* with 1.89 SNPs per 10 kB. Inversions 2 and 3 are more similar at 0.56 and 0.45, respectively, while inversion 1 has only 0.21 SNPs per 10 kB.

We also investigated heterozygous structural variants. We found that there were 90% more duplication, but 22-53% fewer inversion, deletion, and insertion events in the *t* vs *w* part of chromosome 17 (**Figure 3 A-C**). This pattern is different for the total size of these events, where *t* contains a 57% larger summed amount of heterozygous deletions, 70% less deletions, 5% less insertions, and comparatively almost no heterozygous inversions. This is because one inversion in the *w* region is particularly large as it implies essentially the entirety of the *w* region to be heterozygously inverted.

**Figure 3:**
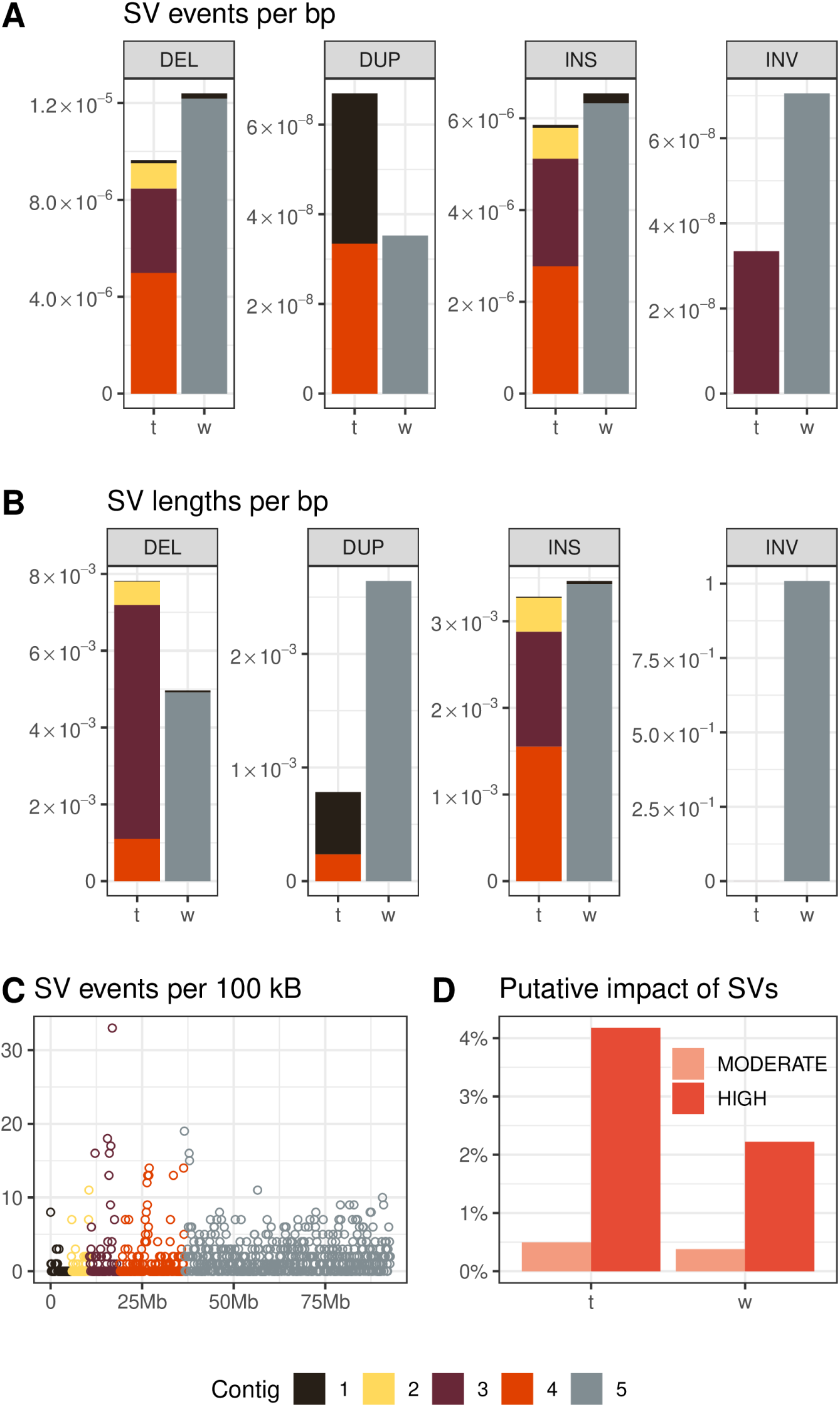
Heterozygous structural variants between GRABS and ILL per non-N base pair in the *t* and *w* regions of chromosome 17. Contig coloring is based on the beginning of each variant. **A)** Events per base pair. **B)** Total summed length per base pair. **C)** *SnpEff* estimated impact of heterozygous structural variants (excluding the lowest possible (non-)impact) in *t* and *w* regions in percent of variants. **D)** Per 100 kB window.

#### Impact of variants

We classified the SNPs heterozygous between the *t* and between the *w*regions of the source populations, which we detected using the *t^wIll^/t^wGrabs^* PacBio long reads, into categories of their putative impact as inferred by *SnpEff*. This revealed a strong difference in the average effect of SNPs in *t* and *w* regions. The estimated effect of SNPs was 3.4 times as likely to be “low,” 4.3 times as likely to be “moderate,” and 3.1 times as likely to be “high” in *t* vs. *w* regions (**Figure 2 F**, **SI Figure 2**). The difference is so large that despite fewer SNPs per base in the *t*, the *t* has absolutely more predicted-to-be-impactful SNPs per base than *w* (1.7*^−^*^5^ vs 1.0*^−^*^5^). The same direction of difference can also be seen in heterozygous structural variants, but to a less extreme degree (**Figure 3 D**, **SI Figure 3**).

#### Candidate genes with putative high-impact differences

The most impactful heterozygous differences between the two *t* variants could be related to the difference in recessive lethality. To provide a list of candidate genes, we filtered the annotated genes to only include at least almost complete transcripts (*≥* 90% amino-acid identity) and compared whether the local sequence where the mutation or deletion occurred is also part of the exon in the reference chromosome 17. This led to a list of 9 *t* genes with missense mutations and 1 gene with a disruption due to a significant deletion (*Fpr2*, Bnip1, Cdkn1a, Cmtr1, Dnah8, Fgd2, Kctd20, Rnf8, Tedc2* and *Vmn2r113*). None of the mutations were found in the list of known mutations on Ensembl.

Some genes stand out as potentially important for phenotypic differences between the *t* variants (**SI Table 2**). *Dnah8* is a dynein subunit that has been proposed to be involved in *t* drive (Fossella et al. 2000; Pilder 2012). *Fgd2* has been found to be important for the *t*’s transmission distortion (Bauer et al. 2007). *Cmtr1* is embryonic lethal in homozygous knockouts at day 8.5 (Lee et al. 2020; Dohnalkova et al. 2023), which is the same time frame as the onset of *t^wIll^*homozygous lethality (Sutter and Lindholm 2015), making it a candidate for the causal variant. *Tedc2* also has reported knockout lethality in utero (Birling et al. 2021). *Rnf8* has a missense heterozygous mutational difference and male mice are known to have impaired spermatogenesis when this gene is knocked out (Li et al. 2010). *t*/*t* male mice are infertile in *t* variants that are not homozygous lethal, thus selection pressure may have been fully absent in this gene since *t*’s inception.

### Differences to the reference chromosome 17

We also compared the assembly chromosome 17 directly with the reference house mouse chromosome 17. The distance to the reference was increased in the *t* region compared to the *w* region with 2.26 times as many SNPs per known base. This is in stark contrast to the decreased heterozygosity in the *t* region (**Figure 4 A** & **C** vs. **Figure 2 A-B**). All tested SVs vs. the reference were also over-represented on the *t* region vs. the *w* region (**Figure 4 B**), between 20% for tandem repeats and 260% for highly diverged regions (and orders of magnitude for inversions).

**Figure 4:**
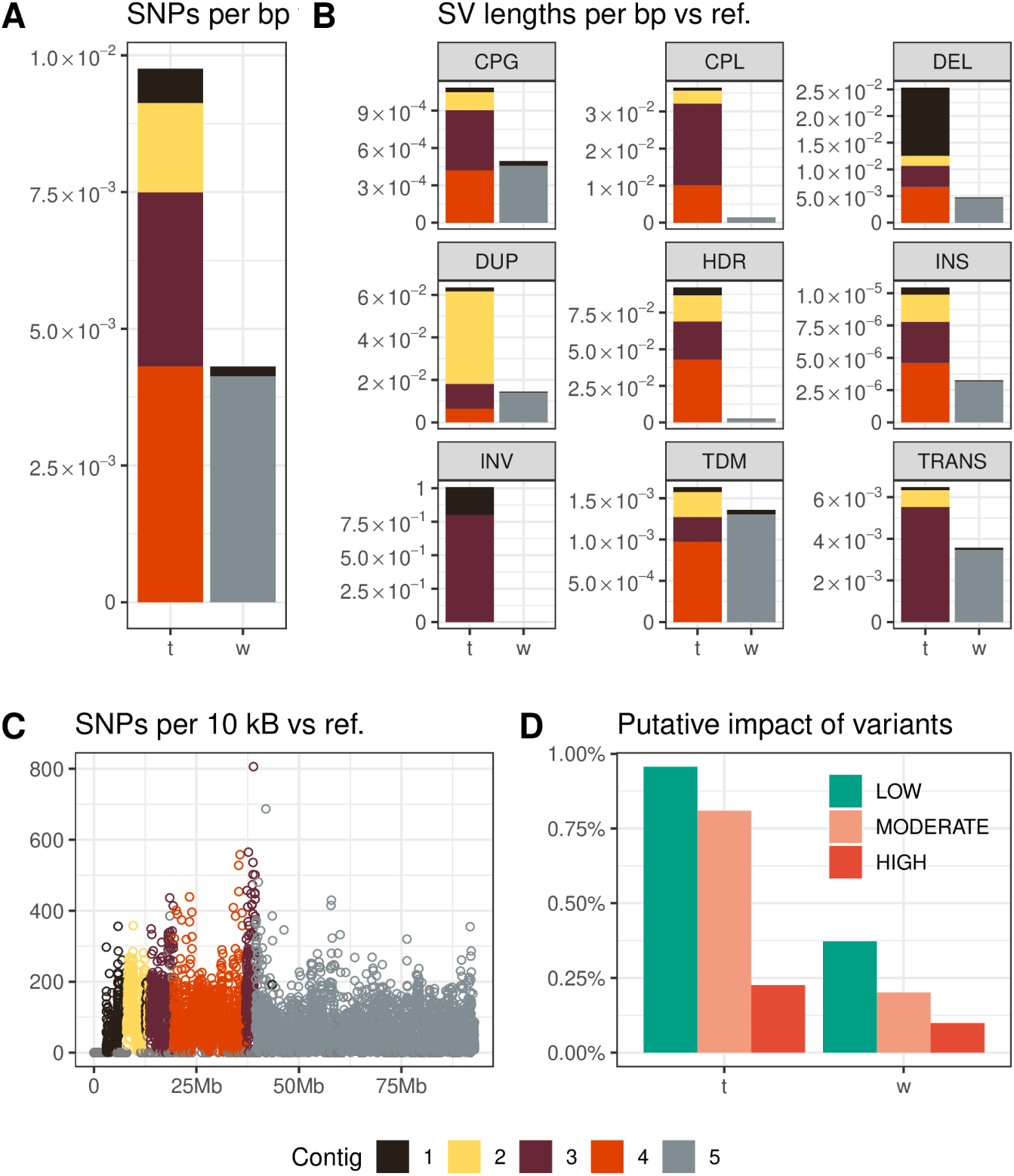
Differences from the reference house mouse chromosome 17 in the aligned regions. **A)** SNPs per base. **B)** Summed lengths of structural variants per base. Contig coloring is based on beginning of each variant. **C)** SNPs per 10 kB. Position is based on the reference. **D)** *SnpEff* estimated impact of reference-aligned variants (excluding the lowest possible (non-)impact) in *t* and *w* regions in percent of variants (both SNPs and SVs).

#### Impact of variants

We also analyzed the estimated impact of the differences to the reference house mouse chromosome using *SnpEff* (**Figure 4 D**, **SI Figure 4**). We found that of all variants (compared to the reference genome), variants in the *t* region were 2.3 times more often of high, 4 times more often of moderate, and 2.3 times more often of low impact (the lowest or likely no impact called “modifier” being thus more common in *w*), which is similar to the heterozygous variants.

### Repetitive elements

To investigate the presence of repetitive elements throughout chromosome 17, we first analyzed the read depth of the Illumina reads across the chromosome. Regions with increased read depth can generally indicate the presence of not fully assembled repetitive DNA, although we cannot rule out some of the changes in read depth being caused by copy number variants. We defined high read depth as being more than twice as high as the average read depth on the chromosome. The *t* region harbored almost two times as many 10 kB windows with high read depth (3.8%) as the *w* region (2%).

In a second step, we used *RepeatMasker* to annotate the assembly’s repetitive elements content. Overall, the fraction of non-N base pairs that is found to be in repetitive elements is very similar between the *t* and *w* region: 42.9% for *t* and 40.5% for *w* (**Figure 5**). These metrics are remarkably similar for the non-*t*-carrying reference house mouse chromosome 17 in the homologous regions: 42.3% and 40.8%. The distribution of classes of annotated repetitive elements is also almost identical to the reference genome (**SI Figure 5**).

**Figure 5:**
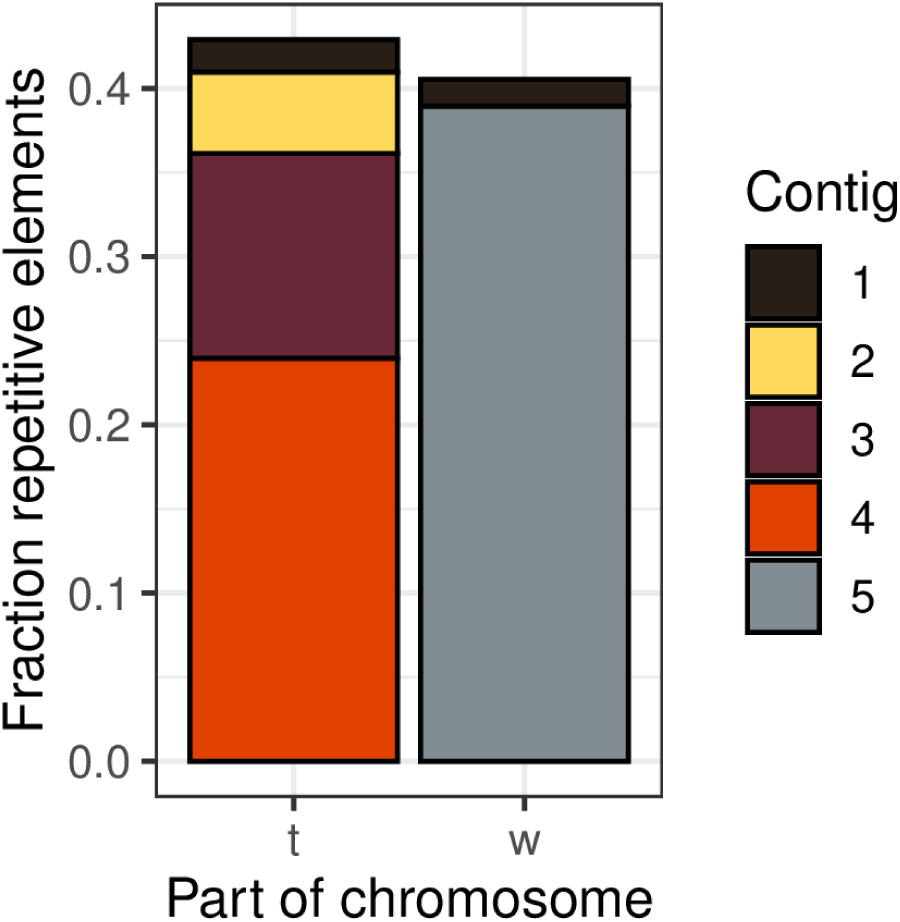
Summed fraction of non-N bases annotated as part of repetitive elements.

### Highly heterozygous region

One region in contig 3 (a *t* region) stood out in both Illumina- and PacBio-based heterozygosity calculations as particularly differentiated between the two *t* variants (**Figure 2 B** & **D**). By searching for consecutive windows with increased heterozygosity in both analyses, we settled on a 1.17 Mb region with an average of 17 (Illumina) to 27 (PacBio) heterozygous SNPs per 10 kB, well above the 1.4 to 3.9 average for *t* or even the 4.6 to 6.4 average for *w*.

The Illumina read depth of this region is average for the contig (6x vs 6.1x), implying usual levels of repetitive elements. However, the annotation by *RepeatMasker* reveals an unusual cumulative size of SINE elements, more than double the *t* region’s average (19% vs. 8%), but overall, repetitive elements only made up 36% of the region’s size, less than in *t* globally.

The estimated effects of SNP variants in the highly heterozygous region are less severe than in its contig overall (“modifier” effect: 98.7% *vs.* 97.1% of variants; “low” effect: 0.8% *vs.* 1.3%; “moderate” effect: 0.5% *vs.* 1.5%; no “high” effects *vs.* 0.05%). However, they are still more severe than in the *w* region.

## Discussion

The evolutionary history and gene content of the *t* haplotype, a widely distributed selfish supergene in the house mouse, has fascinated geneticists for nearly a century. Studying the *t* is challenging, because of high divergence between *t* and *w* combined with the presence of a recessive lethal. We successfully *de novo* assembled the *t* haplotype for the first time using a combination of technologies. This allows us to define the location of the inversions and thereby the coordinates, size, and content of the *t* haplotype. Due to the sampled mouse carrying two different *t* varieties, we were able to compare them for the first time. The *t* region is much more distant to the reference chromosome 17 (from a domesticated strain) than the remaining parts of the assembled chromosome, which are derived from the same wild mice. However, heterozygosity within the *t* region was much lower than heterozygosity in the rest of the chromosome. Hence, even diverged and reproductively isolated strains carry rather similar *t* variants, as predicted by genetic studies of the *t* (Figueroa et al. 1985; Silver et al. 1987; Forejt et al. 1988).

Suppression of recombination by structural variants, such as inversions, allows the integrity of supergenes to be maintained across generations, and is a feature of many supergenes including the *t* haplotype (Black and Shuker 2019). We detailed the four inversions within the *t* haplotype, which was first proposed by Hammer et al. (1989). Distorter loci, that together effect the super-Mendelian inheritance of the *t*, are associated with different inversions (Herrmann and Bauer 2012), and their evolution is not understood. Our assembly had too many unknown bases to directly investigate distorter loci but profiles of SNP heterozygosity suggest an increasing age and/or recombination frequency from inversion 1 to 4. This contrasts with divergence-from-w-based estimates that predict inversion 4 as the youngest inversion (Hammer and Silver 1993; Kelemen and Vicoso 2018). Further, a decreased level of mutation severity in inversion 4 may reflect increased recombination, consistent with findings of Kelemen and Vicoso (2018). More work is needed to untangle these complex results.

Recombination suppression is predicted to result in deterioration, such as an accumulation of deleterious alleles. Both *t^wIll^*and *t^wGrabs^*carry recessive lethal alleles, as homozygotes of each variant die prenatally (Lindholm et al. 2013, Lindholm pers. comm.). Our finding that the two *t* variants have more putatively deleterious differences relative to their size than the much more diverged *w* region of the chromosome fits well with our evidence that *t^wIll^*/*t^wGrabs^* individuals are viable, and therefore do not share the same lethality allele. The large quantity of putatively impactful differences could point towards the presence of more than one lethal allele per *t* variant (Howell et al. 2004), or a complex, multi-allelic base for the different lethalities. We searched for the most promising candidate genes for highly impactful, perhaps lethal, differences and were able to provide 9 genes that are likely significantly affected by heterozygous differences, some of which have known knockout lethality effects associated with them. Although there are numerous different *t*-associated lethal loci (Silver 1985), only two have yet been identified (Sugimoto et al. 2012; Lange et al. 2017). Lethal loci associated with individual variants are important markers of evolutionary history that directly impact fitness, and are also of interest for their window into dysfunctional development.

Many phenotypic traits of the *t* have been uncovered, particularly the ability of the *t* to gain a fertilization advantage by damaging rival *w* sperm during development (Herrmann et al. 1999). Studies on *t^wIll^*-carriers have found a 90% rate of transmission of *t^wIll^*from father to offspring [Lindholm et al. (2013);Sutter:2015caba], increased female longevity (Manser et al. 2011), decreased resting metabolic rates (Lopes and Lindholm 2020), decreased sperm motility (Sutter and Lindholm 2016), altered sperm morphology (Winkler and Lindholm 2022) and poor sperm competitive ability (Sutter and Lindholm 2015), as well as increases in dispersal from their natal population (Runge and Lindholm 2018; Runge and Lindholm 2021; Runge et al. 2022). Other variants of the *t* have been associated with mate choice against *t*-carriers by wildtype individuals (Lenington et al. 1992), changes in territoriality/dominance (Carroll et al. 2004) and trappability (Drickamer et al. 1995). These traits, in particular the increased dispersal, which appears to provide a fitness advantage to the carrier (Runge et al. 2022), demonstrate that the *t* supergene links favorable phenotypes together. Hence, some impactful differences to the reference chromosome, and perhaps between *t*, are likely adaptive, and future work on uncovering causal genes will likely benefit from our assembly as a resource.

However, the great number of impactful mutations, especially compared to the recombining rest of the chromosome, which should be under much more efficient selection, is consistent with deterioration and inefficient purging rather than selection. In contrast, we have no evidence for an expansion of repetitive elements or change in size of the *t* overall, which would also be hallmarks of degeneration. These results fit with the literature’s mixed evidence regarding supergene deterioration (Graves 2006; Svedberg et al. 2018; Stolle et al. 2019; Hill et al. 2022; Komata et al. 2022; Stenløkk et al. 2022). The *t*’s low rates of recombination are expected to predispose it for degeneration (Charlesworth and Charlesworth 2000), but it should be noted that all known wild *t* variants are at least male infertile in the homozygous state (Silver 1985). Some have argued that homozygous lethal *t* variants may be selected over homozygous infertile ones (Charlesworth 1994; Munasinghe and Brandvain 2023), though we have found male infertile *t* to perform much better than lethal *t* in our simulations (Runge et al. 2022). Nevertheless, it is remarkable that the diversity in *t* is so small between reproductively isolated populations and yet so impactful. This implies a more recent ancestor of the *t* than the *w* region. It fits the picture of the *t* as a fast spreader through populations through increased dispersal of its carrier mice, with large fitness gains when entering new populations in which *t* frequencies are low (Runge and Lindholm 2018; Runge et al. 2022). It will be interesting to see more *t* variants mapped against this *t* assembly for a more global comparison.

We hope that our contribution can enable further research into the evolution of the *t* haplotype by mapping other *t* variants to the assembly. Is the pattern of small but meaningful diversity between *t* true more broadly? What are the functional consequences of diversity between *t*, and between *t* and *w*? One could speculate that the *t* is primarily selected through its meiotic drive mechanism and deterioration elsewhere is overcome via rescue by the homologous chromosome’s alleles. This could be investigated by comparing viable variants (in which homozygotes are male-sterile but female fertile) with lethal variants. What can we learn about the diversity of the *t* haplotype within populations? Do they exhibit variation in deterioration? Can we find more evidence and explanations of heterozygous regions within the *t*? Where are different variants and inversions placed in the history of the *t*? To what extent does the genomic structure and evolutionary history of the *t* resemble that of other supergenes or selfish elements?

Our work has revealed the deterioration and content of this selfish supergene, and further comparative studies will help us understand its history.

## Methods

### Study animals

House mice (*Mus musculus domesticus*) of the Alin:ILL (MGI:7579052) and Alin:GRABS (MGI:7579028) wild-derived strains were studied. The ILL strain was founded by wild-caught house mice from the University of Zurich house mouse study population near Illnau-Effretikon, Kanton Zurich, Switzerland (König and Lindholm 2012). The *t* haplotype that occurred in this population (*t^wIll^*, MGI:7579053) has been extensively studied (e.g. Lindholm et al. (2013); Manser et al. (2011); Manser et al. (2020); Runge and Lindholm (2018); Sutter and Lindholm (2015)), but its genetic similarity to previously described *t* variants is unknown. The GRABS strain was founded by wild-caught mice from locations near Grabs, St. Gallen, Switzerland, which carried a different *t* haplotype variant, *t^wGrabs^* (MGI:7579025). Homozygotes of *t^wIll^* die prenatally, as do homozygotes of *t^wGrabs^*, but they complement each other so that *t^wIll^*/*t^wGrabs^* individuals survive. We used a hybrid adult individual for this study, (ILL-*t^wIll^* X GRABS-*t^wGrabs^*)F1. All male hybrid offspring of ILL and GRABS are sterile, without sperm, whether or not they carry *t* haplotypes (Grize et al. 2019). This is because of severe incompatibility of Robertsonian fusions between the populations. F1 hybrids of the two strains form a multivalent chain of 15 chromosomes at meiosis. The karyotype of the ILL strain is designated CHHN with 2n = 24 (1.3 2.8 4.12 5.7 6.15 9.14 10.11 13.16, with the full stop indicating chromosome arms that are fused as a metacentric chromosome) and that of GRABS is CHBU with 2n = 22 (1.18 2.5 3.6 4.12 7.15 8.16 9.14 10.17 11.13; Grize et al. (2019)). We determined that animals carried 0, 1, or 2 copies of the *t* haplotype by PCR polymorphism at the *Hba.ps4* locus (Schimenti and Hammer 1990).

### Sample preparation and sequencing of ***t^wIll^***/***t^wIll^*** homozygous embryos

Resorption of embryos of the genotype *t^wIll^*/*t^wIll^* is underway at 9 days post-copulation (Sutter and Lindholm 2015). We therefore removed embryos at 8 days post-copulation. After phenol-chloroform DNA isolation, we pooled DNA from 6 *t^wIll^*/*t^wIll^* embryos. The Nextera DNA Library Preparation Kit was used for library preparation before paired end (2 x 250bp) sequencing on the Illumina MiSeq platform at the Max-Planck-Institute for Evolutionary Biology using the MiSeq Reagent Kit v.2 500 cycles in 2013. The sequencing data yielded an assembly contigs coverage of 27x due to outlier regions (25th read depth percentile was 3.0, 75th was 10.3).

### Sample preparation, sequencing, and optical imaging from a t*^wIll^*/*^twGrabs^* adult

We extracted 30 µg high molecular weight DNA from 20 mg of liver tissue of the ILL-*t^wIll^*X GRABS-*t^wGrabs^* mouse using Qiagen GenomicTip-20 with the manufacturer’s protocol. Sequencing was done using the with PacBio Sequel. The coverage of the assembly was 39.4x with a 25th percentile read depth of 23.5 and a 75th percentile of 33.1, and an average read length of 7,902 *±* 9,700 (SD).

Similarly, 70 mg liver tissue of the same mouse was used for preparation with the official bionano protocol and the optical imaging was performed on the bionano saphyr platform. We received 7,428,309 molecules with an N50 of 174 Kb and a label density of 14.3 per 100 Kb.

### PacBio *de novo* assembly

We used *wtdbg2* 2.5 (followed by *wtpoa-cns*) with the settings -x sq -g 2.7g to *de novo* assemble the genome based on PacBio long reads (Ruan and Li 2020). QC metrics can be found in Table 2.

**Table 2:**
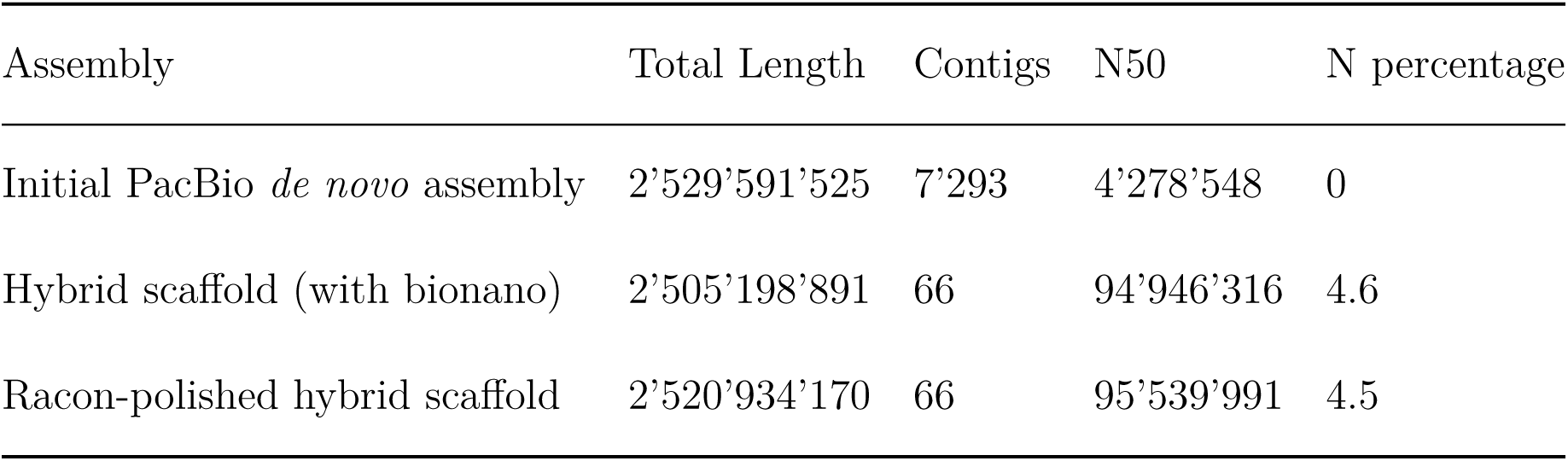
Overview of the *de novo* assembly quality metrics throughout the process.

### Hybrid scaffolding

To increase the assembly quality, we combined the initial long-read-based *de novo* assembly with an optical-imaging-based *de novo* assembly (without reference genome, expected genome size of 2.7 Gb, without trimming) using bionano’s hybrid scaffolder software on bionano Solve 3.6.1. This much improved the assembly (QC metrics can be found in Table 2).

### Polishing

We polished the hybrid scaffolds using Racon 1.5.0 (Vaser et al. 2017) using alignments of the long reads (aligned with minimap2 2.24 (Li 2018) using -x map-pb). QC metrics can be found in Table 2.

### Chromosome 17 contigs discovery

We used *RagTag* 1.0.1 (Alonge et al. 2022) to find the corresponding reference chromosomes for the 66 scaffolds of our assembly. Five contigs were detected as being similar to the reference chromosome 17, the *t*-carrying chromosome, while no contigs matched any unplaced chromosome 17 reference contigs. A further 6 contigs could not be placed, making up 1,358,472 bp in total. All autosomes + X had contigs assigned to them.

### Short-read-based polishing

Finally, we corrected the chromosome 17 assembly using *Pilon* 1.23 (Walker et al. 2014) with *t^wIll^/t^wIll^* homozygous Illumina reads. To speed up the process, we reduced the alignment to only include reads mapping to the chromosome 17 contigs identified before. *Pilon* made 122,670 changes (27,353 single-base additions, 51,154 single base changes, 22,317 single base removals, 4,689 multi-base additions, 16,747 multi-base removals). Changes made by Pilon were also analyzed to interpret the differences between the two *t* variants, as the assembly is based on two variants, so some parts of the assembly will represent one or the other variant.

### Orientation

While *RagTag* can be used to orient contigs according to the reference orientation, but to infer large inversions, such as those that are likely present in *t*, we needed to extract additional information from alignments to infer contig orientation.

For short-reads, we searched for high quality mapped (MQ>=30), non-duplicate, notproperly-paired (i.e. across contigs) reads (-F 1538). Then, we only looked at read pairs where both mates were mapped near the ends (within 500 kB) of different contigs. Such events were taken as evidence for both contigs being close to each other in this particular orientation, for example if one mate mapped to the end of contig 1 and another to the beginning of contig 2, then these contigs are likely in this order and orientation.

We also used the long reads for further evidence. Here, we looked for reads that mapped well (MQ *≥* 30) to ends of different contigs alternatively (i.e. supplementary alignments). We interpreted such reads to support that the two contigs belonged together in that orientation.

### Genome annotation using GeMoMa

We annotated the genome using GeMoMa 1.9 (Keilwagen et al. 2019) with the GeMoMaPipeline options GeMoMa.Score=ReAlign AnnotationFinalizer.r=NO using the mm10 reference genome (GRCm38.p6), followed by result filtering using GeMoMa GAF.

### Differences between the two *t* variants

We used two approaches to quantify the differences between the two *t* variants. First, we analyzed single base changes made by Pilon (see **Short-read-based polishing**). We quantified these changes for each contig in relation to the non-N size of the contig and aggregated those values for the *t* and *w* regions.

Secondly, we re-aligned the PacBio long reads of the heterozygous *t^wIll^/t^wGrabs^* mouse against the final (re-oriented) chromosome 17 assembly. We used *minimap2* 2.24 with -x map-pb for alignment, filtered using *samtools* 1.16.1 (Danecek et al. 2021) for MQ *≥* 30, removed non-primary alignments, used *longshot* 0.4.5 (Edge and Bansal 2019) for calling variants, and filtered those variants for GQ *≥* 30, AC *≥* 10, heterozygosity, and for having at least 80% of alleles be reference or alternative allele (i.e. not read errors). These filtered high-quality heterozygous calls were then quantified per contig in relation to the non-N size of the contig and aggregated those values for the *t* and *w* regions.

Finally, we also quantified heterozygous structural variants. We called structural variants using *Sniffles2* 2.0.7 (Smolka et al. 2024) on the same alignments mentioned for heterozygous SNPs. These results were also filtered for heterozygosity, GQ *≥* 30, and AC *≥* 10. These filtered high-quality heterozygous calls were then quantified per type (e.g. deletions) in their occurrence and also in their total summed length per contig in relation to the non-N size of the contig and aggregated those values for the *t* and *w* regions.

### Impact

To estimate the average impact of heterozygous differences in the *t* and *w* regions, we used *SnpEff* 5.1-2 (Cingolani et al. 2012). We used the GeMoMa assembly annotations to build a custom database for *SnpEff* to refer to (SnpEff build -gtf22 -v tt.Chr17 -noCheckCds -noCheckProtein) and then let *SnpEff* run on both SNP and SV genotypes (independently). We then analyzed all non-modifier (lowest) estimated impact as a fraction of the number of variants, per *t* and *w* region. Often variants have several impact annotations, in that case we only counted them once and only as the highest annotated impact.

#### Candidate genes

To infer possible candidate genes for particularly high impact differences between the two *t* variants, we filtered the results for the highest two impact categories (“MODERATE” and “HIGH”) and only included genes where the assembly was 90% amino-acid identical with the reference. We then blasted the region surrounding each variant (*±* 10 flanking bases) against the reference chromosome 17, testing to see if this sequence is also part of a respective gene’s exon there. These most promising candidates were then examined regarding what is known about them.

### Differences to the reference genome

We used *NUCmer* of *MUMmer* 4.0.0rc1 (Marçais et al. 2018) to align the concatenated assembly contigs against the reference chromosome 17 (*GRCm38.p6*). We ran *SYRI* 1.6 (Goel et al. 2019) to identify structural variants and single nucleotide polymorphisms between the assembly and the reference using a minimum identity of alignments of 80% and a minimum length of 100 bp. We used the *SYRI* base alignments to generate the dotplot.

#### SnpEff variant impact

We ran *SnpEff* very similarly to the heterozygous approach on the variants called by *SYRI* to quantify the putative impacts of mutations and structural changes based on the reference annotation *GRCm38.99*.

#### Repetitive sequences

We annotated repetitive sequences on the assembly and on reference chromosome 17 for comparison using *RepeatMasker* 4.1.5 (Smit, A et al. 2013/2015). We then aggregated the results by class and analyzed the total length of classes of repetitive elements in relation to the non-N size of the *t* and *w* regions.

### Published nucleotide diversity

To compare our levels of heterozygosity with nucleotide diversity in wild house mouse populations, we extracted the levels of nucleotide diversity (*π*) for German, French, and Iranian house mouse populations from the public genome tracks published by Harr et al. (2016) under https://www.user.gwdg.de/~evolbio/evolgen/wildmouse/m_m_domesticus/browser_tracks/pi_bigWig/ and aggregated all windows into mean and standard deviation values.

## Data availability

(to be changed) A jbrowse2 server has been set up at https://janniklasrunge.de/jbrowse2/server/ for easy exploration of the assembly and its associated data. The Racon-polished hybrid scaffold whole genome assembly, as well as the raw Illumina and PacBio data is at SRA BioProject PRJNA1104182. The final, Pilon-polished and oriented chromosome 17 assembly (with annotations) we presented here and PacBio alignments, as well as bionano molecule data can be accessed at Zenodo under 10.5281/zenodo.11066354.

## Acknowledgements

We thank Diethard Tautz, Andrés Bendesky, and Barbara König for helpful discussions at several stages of this project, and computing support. We are grateful to Barbara König for leading the Illnau barn study, establishing the ILL laboratory strain, and providing infrastructure support, Sofia Grize for establishing the GRABS strain, Andreas Sutter for isolating *t*/*t* embryos, and Jari Garbely and Rie Shimizu-Inatsugi for DNA extraction. We also thank Sven Künzel for performing library preparation and Illumina sequencing at the Max Planck Institute for Evolutionary Biology, and Lucy Poveda and Anna Bratus of the Functional Genomics Centre Zurich for sequencing and providing support for PacBio and BioNano platforms. Funding was provided by the Swiss National Science Foundation grants 138389, 160328, and 189145 and from a pilot project grant from the University Research Priority Program Evolution in Action.

## Permits

The work was approved by the Veterinäramt of the Kanton Zurich under permits 110/2013 and 64/2014.

