## Supplementary material for "*De novo* assembly of the selfish *t* supergene reveals a deleterious evolutionary trajectory": SI Figures and Tables

September 15, 2024

[1] Max-Planck Institute for Evolutionary Biology, Plön, Germany

[2] Department of Evolutionary Biology and Environmental Studies, University of Zurich, Zürich, Switzerland

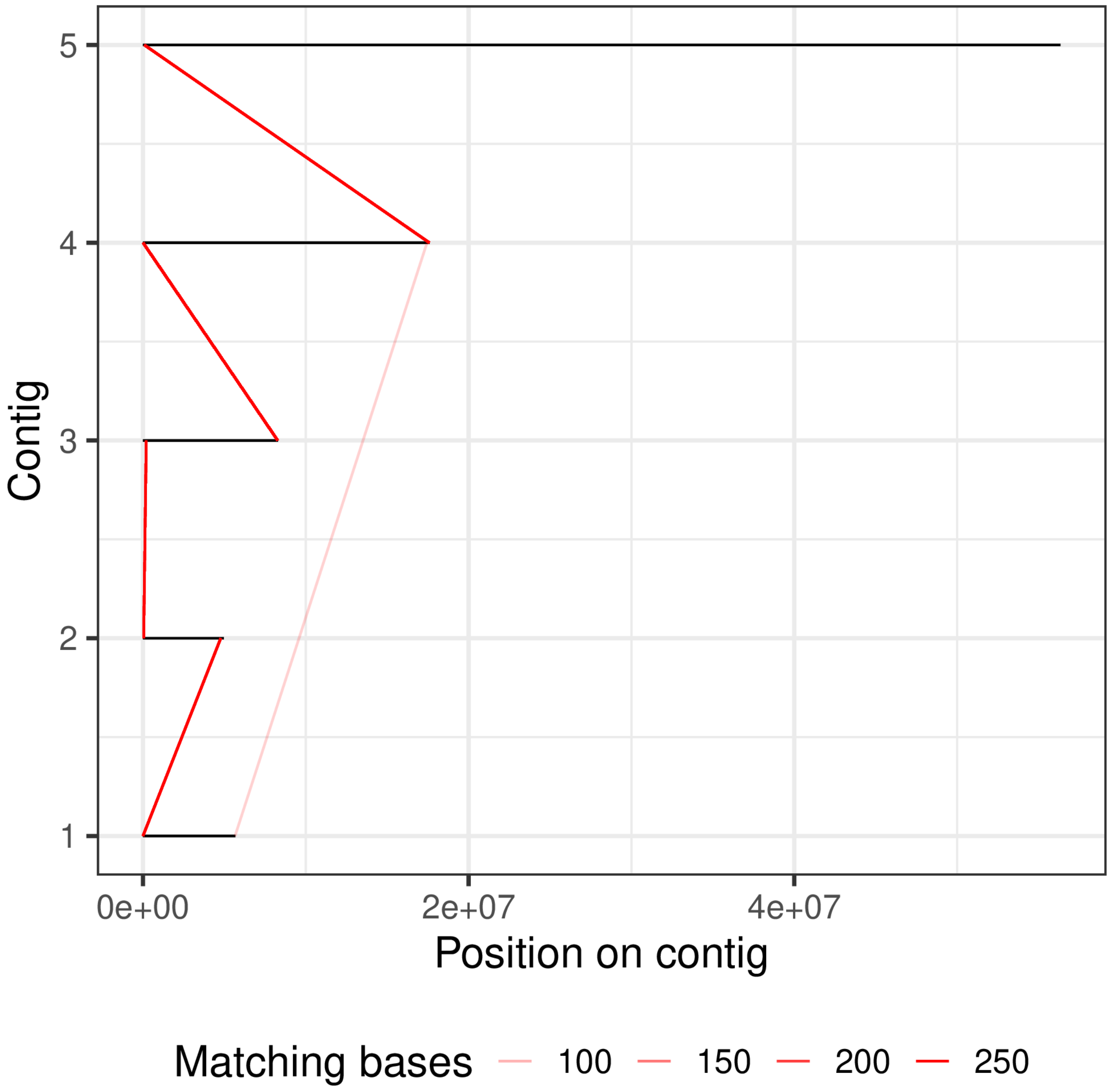


**Figure 1:** Illumina paired-end reads that connect the contigs in their original orientation near the ends of each contig (excluding all others).

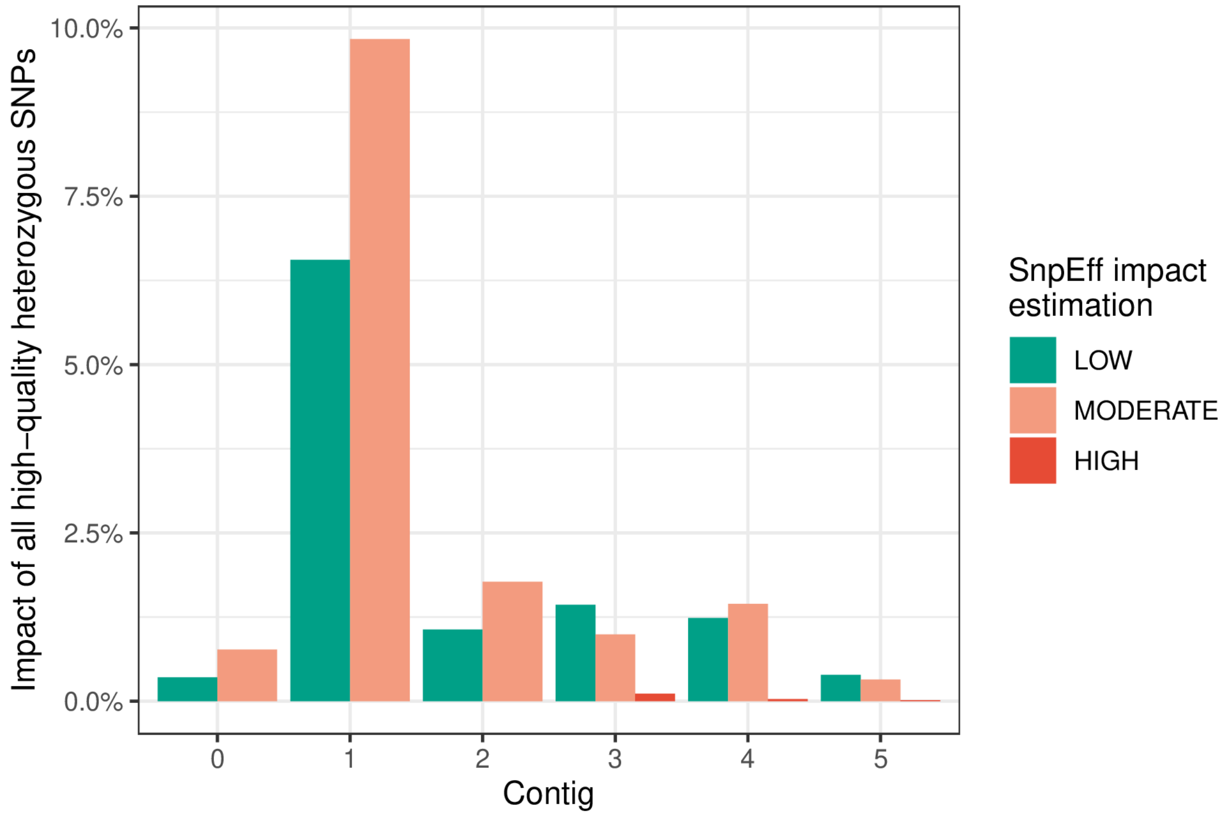


**Figure 2:** Overview of the frequency of higher-than-modifier putative impact of high quality heterozygous SNPs in the ILL x GRABS mouse. Contig 0 is the first $w$ part of contig 1, so contigs 0 and 5 are $w$ regions, and 1 to 4 are $t$ regions. This is the same data as in (**MS Figure 2 F**)

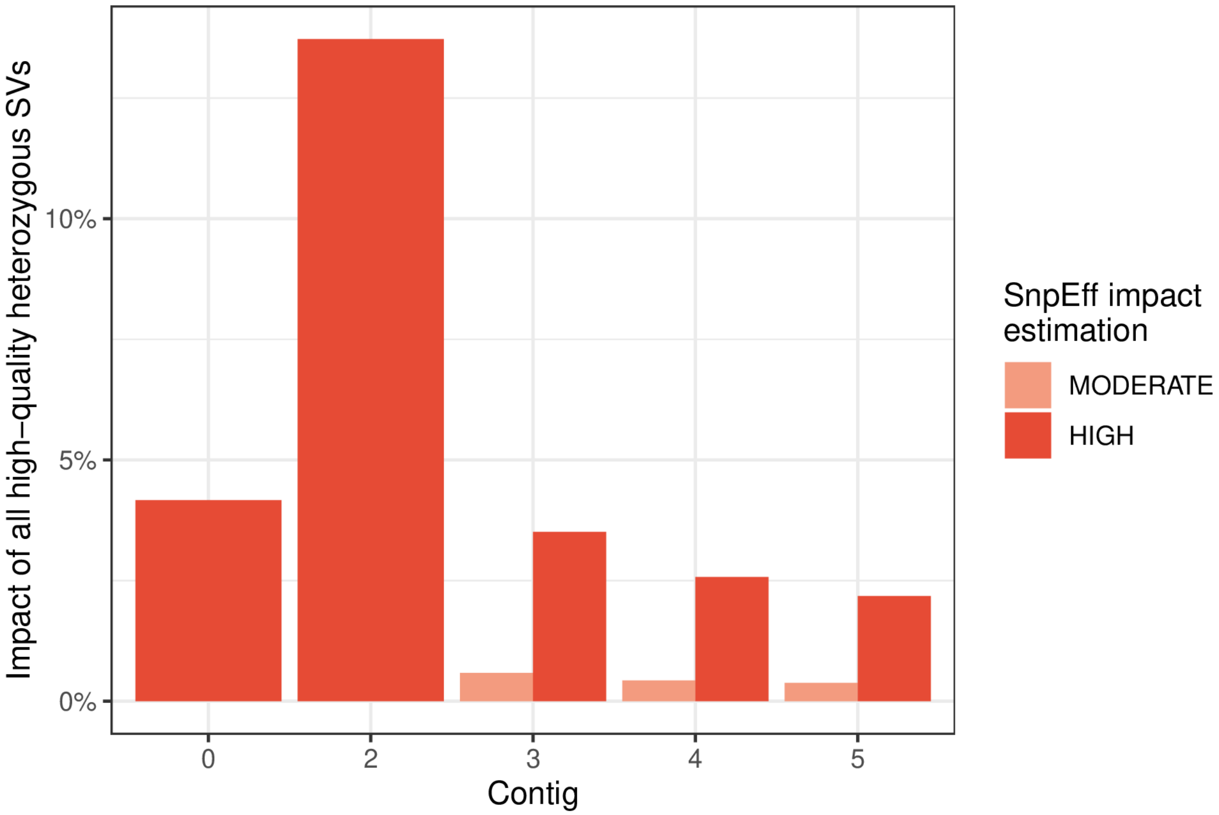


**Figure 3:** Overview of the frequency of higher-than-modifier putative impact of high quality heterozygous SVs in the ILL x GRABS mouse. Contig 0 is the first $w$ part of contig 1, so contigs 0 and 5 are $w$ regions, and 1 to 4 are $t$ regions, with 1 being absent for lack of higher-than-modifier impact SVs. This is the same data as in (**MS Figure 3 D**)

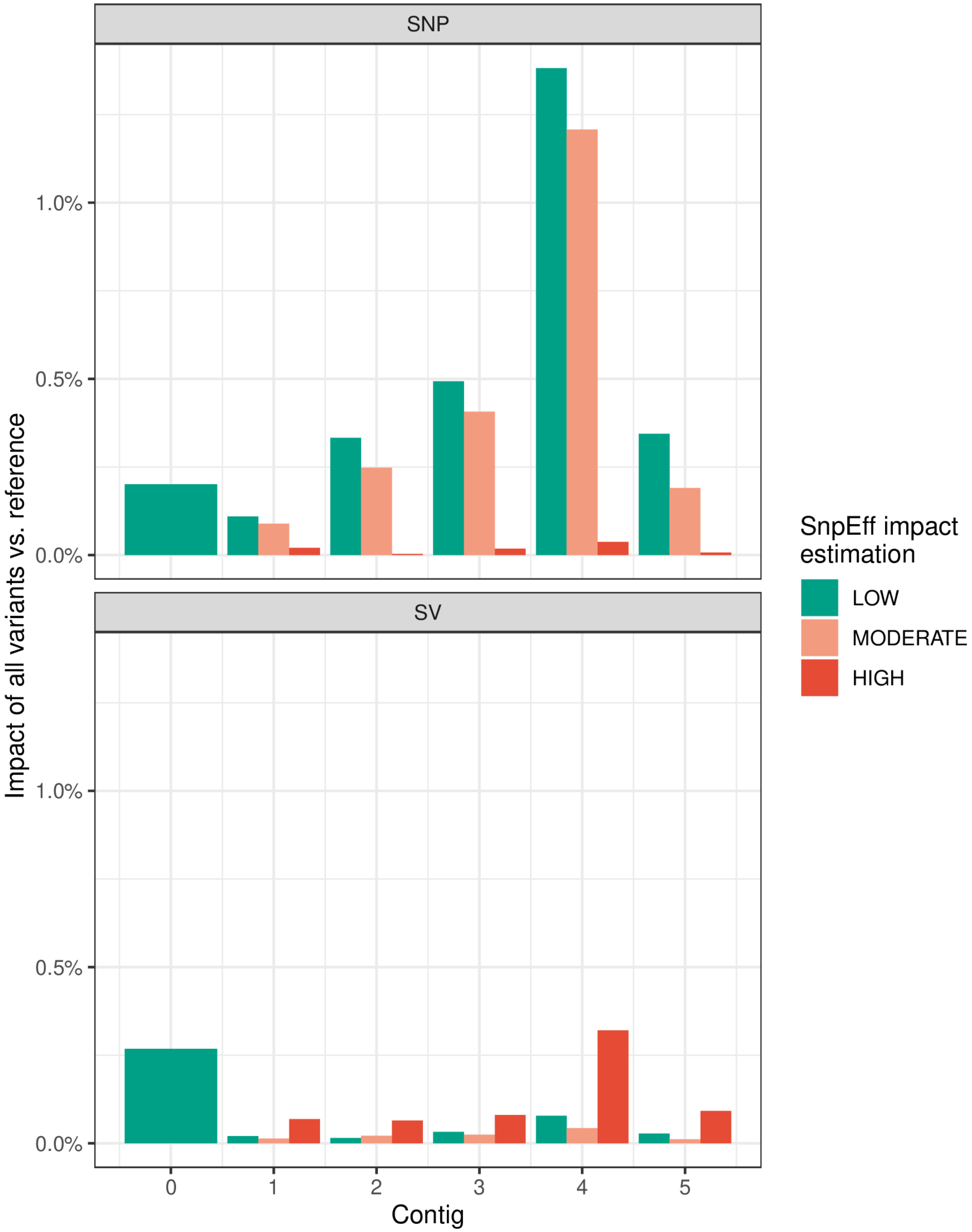


**Figure 4:** Overview of the frequency of higher-than-modifier putative impact of chromosome 17 assembly-vs-reference variants, separated into SNP and SV. Contig 0 is the first $w$ part of contig 1, so contigs 0 and 5 are $w$ regions, and 1 to 4 are $t$ regions. This is the same data as in (**MS Figure 4 D**)

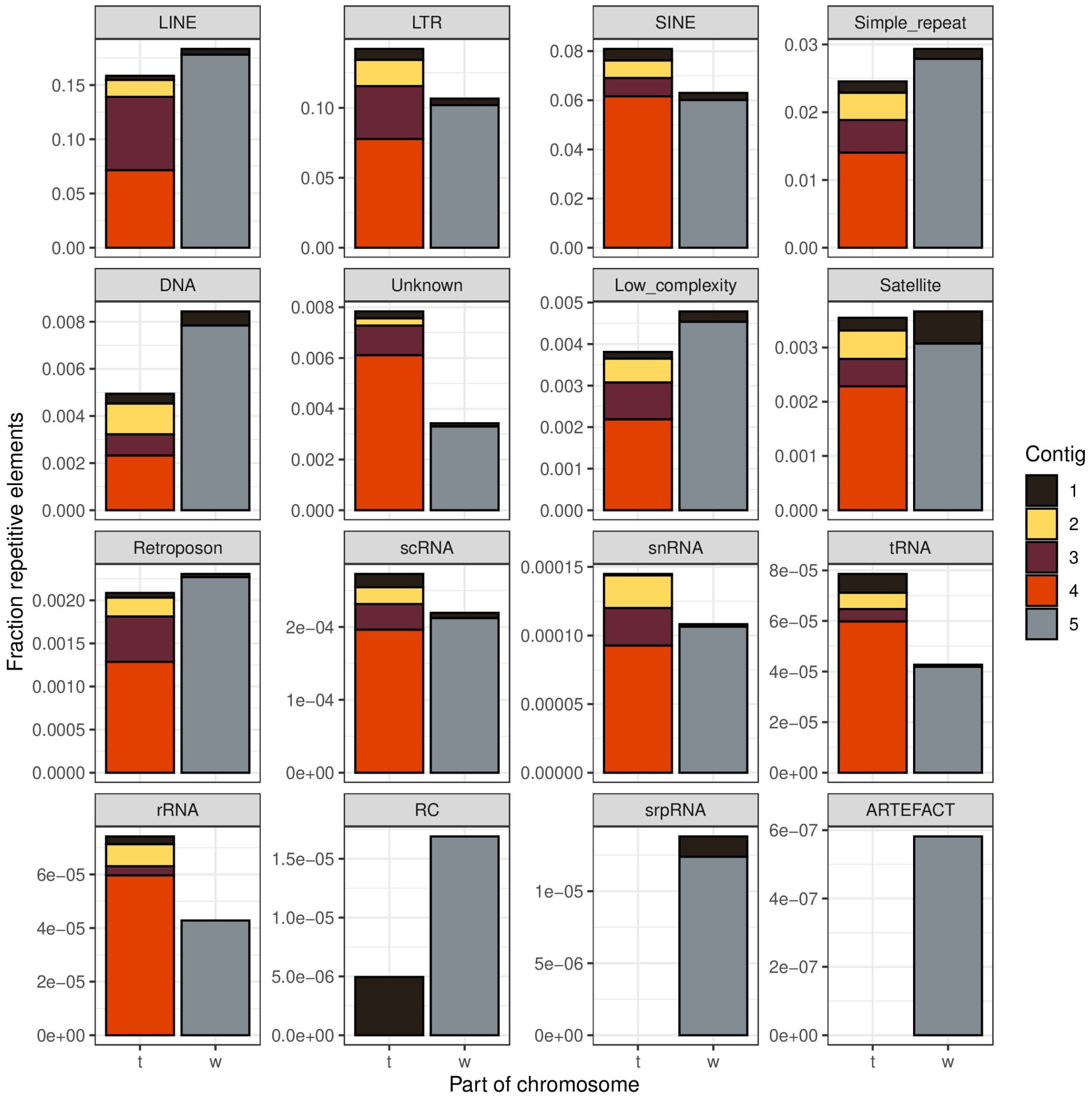


**Figure 5:** Overview of the fraction of DNA that was detected as one of 16 repetitive element families by RepeatMasker in $t$ or $w$ region, colored by each contig's relative contribution to that fraction.

**Table 1:** Overview of the evidence of contig orientation. The chosen order & orientation is given in bold. An asterisk indicates that the contig was flipped in its orientation compared to the initial assembly output.

| Contig connection | Short read support | Long read support |
|---|---|---|
| **1*-2*** | 3 paired-end reads | 60 long reads |
| **2*-3** | 3 paired-end reads | 43 long reads |
| **3-4** | 20 paired-end reads | 594 long reads |
| **4-5** | 5 paired-end reads | 98 long reads |
| 2*-1* | 0 | 174 |
| 4*-1* | 0 | 95 |
| 4-1* | 0 | 64 |
| 5-1* | 0 | 63 |
| 1-2* | 0 | 42 |
| 2*-4 | 0 | 30 |
| 3-1* | 0 | 29 |
| 5*-1* | 0 | 23 |
| 1-3 | 0 | 22 |
| 1*-5 | 0 | 15 |
| 5-1 | 0 | 15 |
| 5*-4 | 0 | 9 |
| 4*-2* | 0 | 9 |
| 4*-3 | 0 | 7 |

| Contig connection | Short read support | Long read support |
|---|---|---|
| 1*-4 | 0 | 7 |
| 5-4 | 0 | 7 |
| 3-2* | 0 | 7 |
| 5-2* | 0 | 7 |
| 2*-5 | 0 | 7 |
| 5-3 | 0 | 7 |
| 1*-2 | 0 | 6 |
| 4-1 | 0 | 5 |
| 3*-2* | 0 | 5 |
| 3-2 | 0 | 5 |
| 5*-2* | 0 | 5 |
| 5-3* | 0 | 5 |
| 4-3 | 0 | 4 |
| 1*-3* | 0 | 4 |
| 3-5 | 0 | 4 |
| 4-3* | 0 | 3 |
| 1*-3 | 0 | 3 |
| 4-2* | 0 | 3 |
| 4-2 | 0 | 3 |

| Contig connection | Short read support | Long read support |
| --- | --- | --- |
| 5-2 | 0 | 3 |
| 5-4* | 0 | 2 |
| 5*-3 | 0 | 1 |

**Table 2:** Candidate genes for large differences between $t$ variants. The assembly coordinates and exact changes can be found in the **SI Tables 3-4** (Excel). "DE" stands for differential expression information from Lindholm *et al.*, 2019. Embryonic expression information is based on Bgee.

| gene | DE in adult $w^{wIll}/t^{wIll}$ vs $w^{wIll}/w^{wIll}$ | Type found | SnpEff details | Embryonic expression of gene |
| --- | --- | --- | --- | --- |
| Fpr2 | Not expressed in testis. Expressed in ovaries, liver, brain | $t$-het. SNP | missense variant | Yes |
| Dnah8 | Reduced expression in testis, ovaries, liver (male); regularly expressed in liver (female) and brain | $t$-het. SNP | missense variant | Yes |
| Cmtr1 | Reduced expression in testis and liver (male); alternative splicing in ovaries and brain, regular expression in liver (female) | $t$-het. SNP | missense variant | Yes |

| gene | DE in adult $w^{wIll}/t^{wIll}$ vs $w^{wIll}/w^{wIll}$ | Type found | SnpEff details | Embryonic expression of gene |
|---|---|---|---|---|
| Rnf8 | Alternative splicing in ovaries and brain (male & female), regular expression in testis and liver | *t*-het. SNP | missense variant | Exclusively |
| Fgd2 | Reduced expression in testis; regularly expressed in ovaries, liver, and brain | 2 *t*-het. SNPs | missense variants | Yes |
| Cdkn1a | Expressed in testis, ovaries, liver, brain | *t*-het. SNP | missense variant | Yes |
| Kctd20 | Increased expression in testis; regularly expressed in ovaries, liver, brain | *t*-het. SNP | missense variant | Exclusively |
| Tedc2 | Expressed in testis, ovaries, liver, brain | *t*-het. SNP | missense variant | Yes |
| Vmn2r113 | Not expressed in testis, ovaries, liver, brain | *t*-het. SNP | missense variant | Unknown |

| gene | DE in adult $w^{wIll}/t^{wIll}$ vs $w^{wIll}/w^{wIll}$ | Type found | SnpEff details | Embryonic expression of gene |
|---|---|---|---|---|
| Bnip1 | Increased expression in testis; regularly expressed in ovaries, liver, brain | $t$-het. deletion | bidirectional gene fusion, frameshift variant, splice region variant & conservative inframe deletion | Yes |

Detailed information on the putatively impactful $t$ heterozygous SNPs / SVs can be found in the SI.xlsx (SI Tables 3-4).
